# Multiplicative Updates for Optimization Problems with Dynamics

**DOI:** 10.1101/234526

**Authors:** Abbas Kazemipour, Behtash Babadi, Min Wu, Kaspar Podgorski, Shaul Druckmann

**Affiliations:** Department of Electrical and Computer Engineering, University of Maryland, College Park, MD, USA; Janelia Research Campus, Ashburn, VA, USA

**Keywords:** NMF, point process smoothing, Poisson image reconstruction, nonnegativity, KKT conditions, latent variables

## Abstract

We consider the problem of optimizing general convex objective functions with nonnegativity constraints. Using the Karush-Kuhn-Tucker (KKT) conditions for the nonnegativity constraints we will derive fast multiplicative update rules for several problems of interest in signal processing, including non-negative deconvolution, point-process smoothing, ML estimation for Poisson Observations, nonnegative least squares and nonnegative matrix factorization (NMF). Our algorithm can also account for temporal and spatial structure and regularization. We will analyze the performance of our algorithm on simultaneously recorded neuronal calcium imaging and electrophysiology data.

## I. Introduction

The advent of big data has given rise to new challenges in signal processing. Fast and scalable solvers for solving large optimization problems remains a big challenge of optimization theory. In this paper we consider the problem of solving general optimization problems under nonnegativity constraints. Such optimization problems arise in many applications of interest. Examples include nonnegative matrix factorization for images of objects [1], Poisson image reconstruction [2], point process smoothing for stimulus-response experiments in neurophysiology [3], nonnegative least squares [4] and nonnegative calcium deconvolution [5]. In this paper we will use the KKT conditions [6] to provide a unified framework for solving such optimization problems with nonnegativity constraints. As we will see these conditions naturally lead to multiplicative updates with suitable convergence in many applications.

Multiplicative updates have been used for solving ML and MAP estimation as well as KL-divergence minimization. Many of these algorithms are special cases of the so-called proximal backward-forward scheme [7]. These algorithms try to find fixed points of a set of equations resulting from setting gradients of the objective function to zero. A With the help of parallel computing and graphics processing units (GPUs), these iterative methods can be solved very fast. Therefore, they become increasingly important. An important application of these multiplicative updates is the Richardson-Lucy (RL) algorithm for image deconvolution [8], which is widely used in astronomy and microscopy [9]. The RL algorithm recovers the ML estimate of a sample under Poisson statistics [10].

Multiuplicative updates are commonly contrasted with gradient descent methods. Their update steps do not necessarily follow the direction of the steepest descent. Multiplicative updates are argued to be insensitive to noise and more flexible [11]. Despite fast early convergenece multiplicative updates are claimed to converge slowly in later stages [12]. However, this argument has been refuted for Poisson Image reconstruction [11], the Weiszfeld problem [7] and NMF [13] by showing their equivalence to a Majorization Minimization (MM) algorithm which has linear convergence in iterations [14]. In contrast, both multiplicative updates and gradient descent based algorithms such as the proximal-gradient method have sublinear rate of convergence [7] in general. Moreover, with specific choices of the stepsize, in many cases such as the Weiszfeld problem these algorithms have proven to be equivalent [7]. These findings suggest that slow convergence of multiplicative updates in some cases is due to absence of strong convexity in the objective function.

An advantage of multiplicative updates over gradient descent based algorithms is their flexibility in terms of adapting to the objective functions without the need for calculation dual functions or tuning extra parameters such as the step-size. Despite the recent breakthroughs in choosing these parameters [15], each step in calculation of the step size is usually as costly as an iteration of the algorithm which is not as effective for big data problems. In addition many problems such as image reconstruction and calcium deconvolution [16] are spatially separable and are easily parallelized.

Finally, temporal dynamics and penalization play an important role in signal recovery from noisy data. Examples include state-space estimations, video reconstruction and total variation denoising problems. Apart from special cases, the solutions to these problems are generally batch mode and computationally demanding. In this paper we provide a unified framework for generalizations of multiplicative updates to the problems with nonnegativity constraints and dynamics by adapting the update rules to different forms of penalties. We have empirically found that multiplicative updates show superior convergence properties and speed to gradient descent methods for models that include dynamics and penalization.

## II. Notations and Problem Formulation

Throughout the paper we will use the following notation. We use the convention [*T*] = {1,…,*T*} and **W**_[*T*]_ = [w_1_,…,w_T_], i.e. w_*k*_ represents the *k*th column of **W**_[*T*]_. ⊙ and ⊘ denote elementwise multiplication and division respectively. Throughout the paper we will use the terms innovations and spikes interchangeably. Unless otherwise stated, a function acts on a vector elementwise. For a matrix **A** = [*a_ij_*] ∊ ℝ^m × n^ its mixed *p, q*-norm is denoted by ║A║_*p,q*_, i.e.

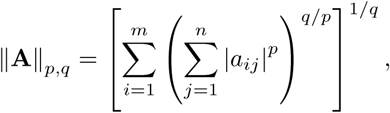

and 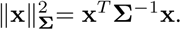 Finally, for a summation

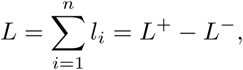

where 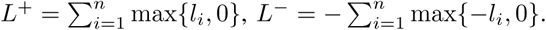

We consider a convex optimization problem of the form

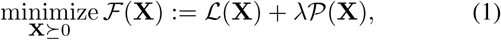

where 𝓛(.) denotes a convex objective function and 𝓟(.) denotes a suitable penalty function. Typically 𝓛(.) is a negative log-likelihood and 𝓟(.) is a smooth norm. Additionally we make the assumption that both 𝓛 and 𝓟 are differentiable with respect to X on the positive orthant,

Among the algorithms used for solving (1) one can name the primal-dual algorithm and proximal gradient method. For specific choices of the penalty functions *𝓁*_1_ and *𝓁*_2_ (Tikhonov) regularization several fast algorithms exist. However these algorithms cannot be easily generalized to arbitrary penalties or temporal dynamics. In some cases such as the gradient based methods they require knowledge of the proximal map or have extra parameters such as the step size to be tuned and chosen. Calculation of the step size is usually as costly as a few iterations of the algorithm and could slow them down. However, our approach to solving (1) does not require tuning of extra parameters and is very simple to implement. We will next discuss our solution.

## III. Solution to the Main Optimization Problem

In this section we will introduce our solution to (1) via multiplicative updates. The Lagrangian form of (1) is given by

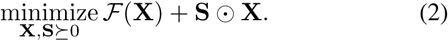

Assuming convexity and zero duality gap, the KKT conditions for (2) can be expressed as

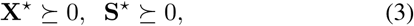

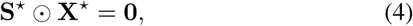

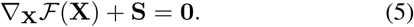

In the rest of the paper, we drop the subscripts and arguments whenever they can be understood from the context. Multiplying (5) by X and using (4) we obtain:

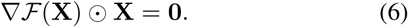

Our solution to (1) looks for a positive fixed point of (6). Therefore giving us the multiplicative update rule

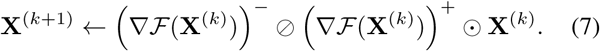

In all application introduced in this paper we initialize the algorithm with a positive solution, the choice of which depends on the application. The update rule will then ensure the solution remains positive. In order to provide more insight into our algorithm we will next provide several examples and applications.

In applications of interest in this paper we consider temporal dynamics in X, hence referring to our algorithm by FAst DEconvolution (FADE) algorithm. In the spirit of easing reproducibility, we have made MATLAB implementations of our codes publicly available [17].

## IV. Examples and Application to Real Data

In this Section we will provide examples of the multiplicative updates in different applications of interest.

### A. Nonnegative Deconvolution

In its simplest form the nonnegative deconvolution problem can be formalized by considering the state-space model given by

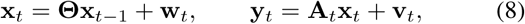

where 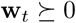 models the innovations at time *t* ∊ [*T*]. Usually, the observation noise is assumed to be i.i.d normal, i.e. v_*t*_ ∼ 𝒩(0, Σ_*t*_) and the measurement matrices **A**_*t*_ are assumed to conserve positivity. For this problem we can identify **W** = **W**_[*T*]_ and

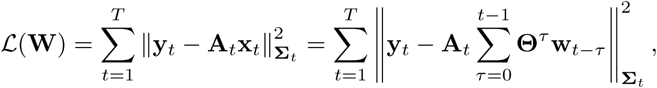

from which we can calculate

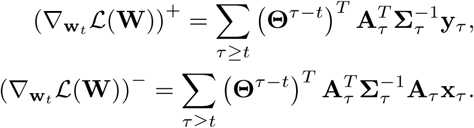

Typically one can use a smooth norm in order to enforce prior assumptions on the spikes, for example one can use a sparsity inducing prior 𝒫 = ║**W**║_1,1_, for which (∇𝒫)^+^ = 1 and (∇𝒫)^−^ = 0. The choice of the penalty function on the spikes is arbitrary and could differ from application to application. In applications where such information is not readily available, one would like to enforce minimal assumptions on the spikes and hence would want to enforce non-informative priors. The most famous example of such priors is known as Jeffrey’s prior [18]. However this problem is an active area of research as there is no unanimously agreed upon choice of non-informative priors.

**Fig. 1:**
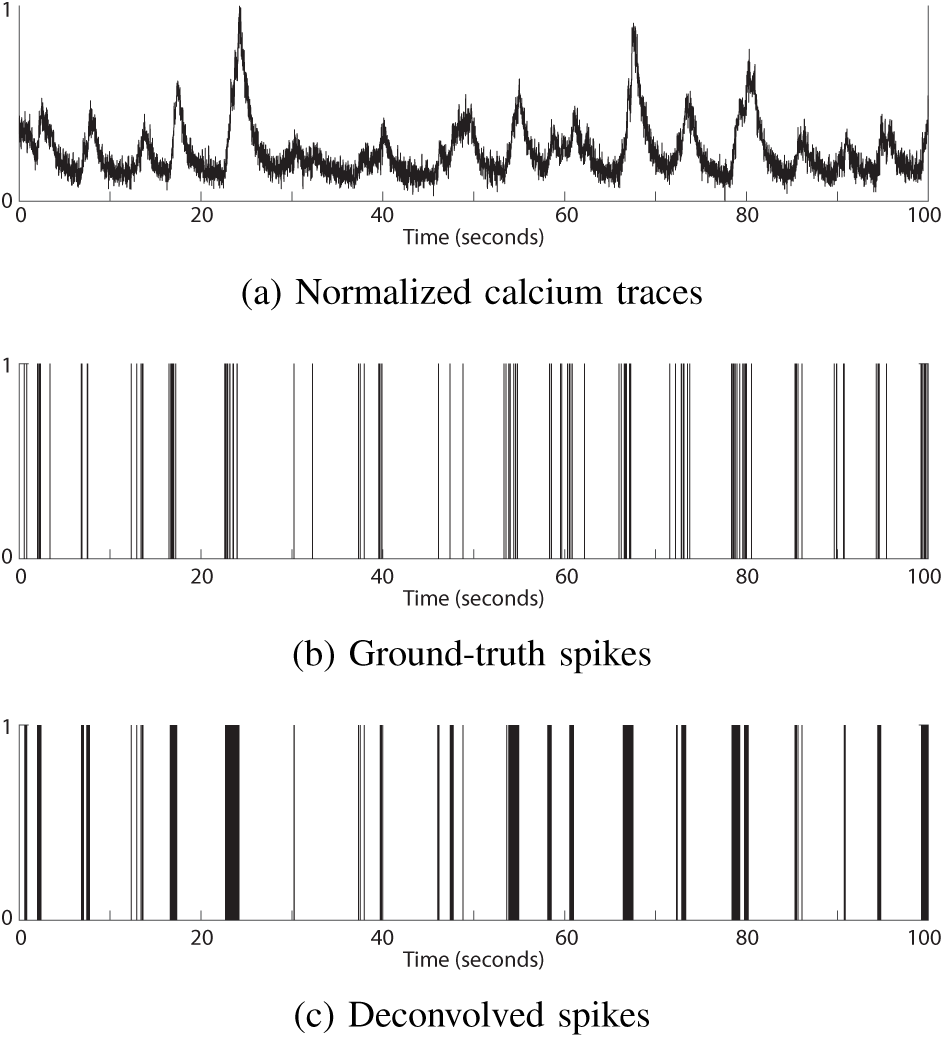
Application of the FADE algorithm to calcium deconvolution problem.

### B. Application to Calcium Deconvolution

Calcium imaging is used to visualize currents associated with action potentials in living neurons. This is done using fluorescent molecules that change their fluorescence properties upon binding calcium, and using a one- or two-photon fluorescence microscope to record these changes [19], [20]. Inferring action potentials (spikes) from calcium recordings, referred to as calcium deconvolution, is an important problem in neural data analysis. For the special case of calcium imaging we have Σ = σ^2^**I**, **A**_*t*_ = **I** and Θ = *θ***I**. Here the baseline is assumed to have been estimated and subtracted separately, but can be estimated similarly. We refer to [5] for details on estimation of the unknown parameters σ^2^ and *θ* and a list of methods used for calcium deconvolution. These approaches require solving convex optimization problems, which do not scale well with the temporal dimension of the data.

Figure 1 shows application of the FADE algorithm to simultaneously recorded imaging and electrophysiology data. The algorithm has covnverged (less than 0.5% change in spikes) in 28 iterations. The data is a 100 second interval from the spikefinder challenge [21] (dataset 3, neuron 1). We have used an AR(2) model and an 𝓁_0.5,1_ penalty on the spikes in order to enforce temporal sparsity. The spikes have been obtained by simply thresholding the deconvolved spikes at 3σ, where σ is the estimated standard deviation of the observation noise. A comparison of the performance of our algorithm with many other methods is provided on the spikefinder challenge website [21].

One can use spatial regularization on elements of w_*t*_ in this setup as well as compressive sensing regimes for when **A** satisfies the restricted isometry property RIP [5]. We refer to [5] for a more detailed discussion.

### C. Poisson Image Reconstruction and Point Process Smoothing

State-space models with Poisson observations have also been studied in many applications of interest. In neuroscience, temporal dynamics of stimulus-response experiments in neurophysiology have been modeled using a Poisson state-space model. In emission tomography, dynamics of the photons hitting the detectors can be modeled with Poisson noise models. Without loss of generality we consider the state-space model given by

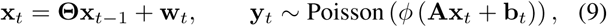

where 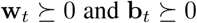 model the spikes and baseline rates at time *t* ∊ [*T*] respectively and *ϕ*(.) is a bijective convex function. Common examples include *ϕ*(*x*) = exp(*x*), *ϕ*(*x*) = 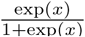 and *ϕ*(*x*) = *x*. We assume the latter in our derivations due to space considerations.

Several approaches have been proposed in the literature for finding the MAP solution to (9). We refer to [22] for a detailed list of these methods. In [3] the authors used the maximum a posteriori derivation of the Kalman filter and proposed an approximate expectation maximization (EM) approach to this problem by Gaussian approximations of the posterior likelihood. This EM approach has several shortcomings. First, it requires solving a nonlinear system of equations which could potentially be computationally costly. Second, it only accounts for Gaussian spikes. Third its performance heavily depends on the Poisson rate model, especially when the rates are small, which is the usual case for spiking activities. In these cases usually *ϕ*(*x*) = exp(*x*) is considered for stability of approximations. Moreover due to nonlinear recursive filtering nature of the problem, the performance of the Gaussian approximation quickly degrades as the dimension of the latent space goes beyond 2 or 3. Similarly, in [22] the authors proposed SPIRAL which uses a Gaussian approximation to *L* and is a gradient-based solution to (9). Except for the special cases of *𝓁_1_* and TV penalties, calculation of the Gaussian model is tedious leading to slow convergence. In [23] the authors introduce a variational auto-encoder (gradient descent based) model to retrieve the low-dimensional temporal factors.

In applications such as fluorescence microscopy, it is also common to use to use variance stabilizing transforms [22] such as square root filtering [24] in order to make Gaussian approximations to the Poisson distribution. In the high photon regime such transformations are not necessary as one can use infinite divisibility property of the Poisson distribution for Gaussian approximations. However one would then need to deal with complications arising from equality of the mean and the covariance matrices for such approximations. In contrast, our algorithm gives an exact solution, is fast, can account for any rate model and suitably scales with the problem dimensions.

The Gaussian approximations could then be used as an input to a Kalman smoother if the innovations (spikes) follow a half-normal or Gaussian distribution. Despite the fact that our solutions are faster, exact and do not involve approximations, for Gaussian state-spaces the Kalman smoother provides a smoothed estimate of the covariances which could be used for building confidence intervals, whereas the covariances are not a direct output of the multiplicative updates.

Considering the MAP estimator for **W** = **W**_[*T*]_ we can identify

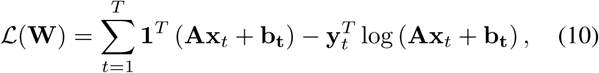

for which we have

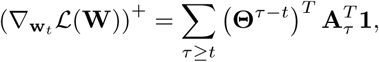

and

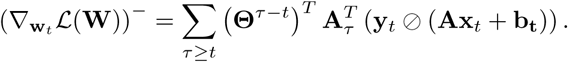

The penalty function and the corresponding terms can be calculated similar to the nonnegative deconvolution problem. A similar update rule can be derived for the baseline. The special case of Θ = 0 (no dynamics with the convention 0^0^ = **I**) and λ = 0 (no penalization) is known as the Richardson-Lucy (RL) iterations. The RL algorithm has also been used with TV seminorm regularization in [25]. Similar to the RL algorithm we can use FADE for blind deconvolution, when the measurement matrix A is unknown. in this setup one can alternatively update A and X. We can also used FADE, for estimation of GLM models for self-exciting point process models [26].

### D. Combination with Other Constraints

In many applications of interest the optimization problem could also include several inequality constraints. For example in fluorescence microscopy the maximum changes of the fluorescence level with respect to baseline (also referred to as 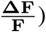 is controlled by the properties of the indicator in use. In these situations we need to satisfy the KKT conditions for the extra constraints. Here we will introduce an adaptive method in order to achieve this goal. Consider the modified problem setup of Section IV-C given by

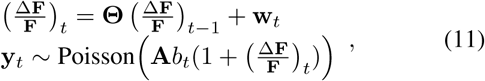

where *b_t_* ≥ 0 denote(the known baseline fluorescence at time *t*, on top of which 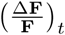 lies. In addition to nonnegativity constraints we need to account for the following constraints

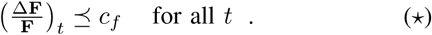

The constant *c_f_* is a characteristic of the indicator used and is assumed to be known. In order to enforce (*) we proceed as in Algorithm 1.

##### Algorithm 1 Multiplicative Updates with Adaptive Regularization

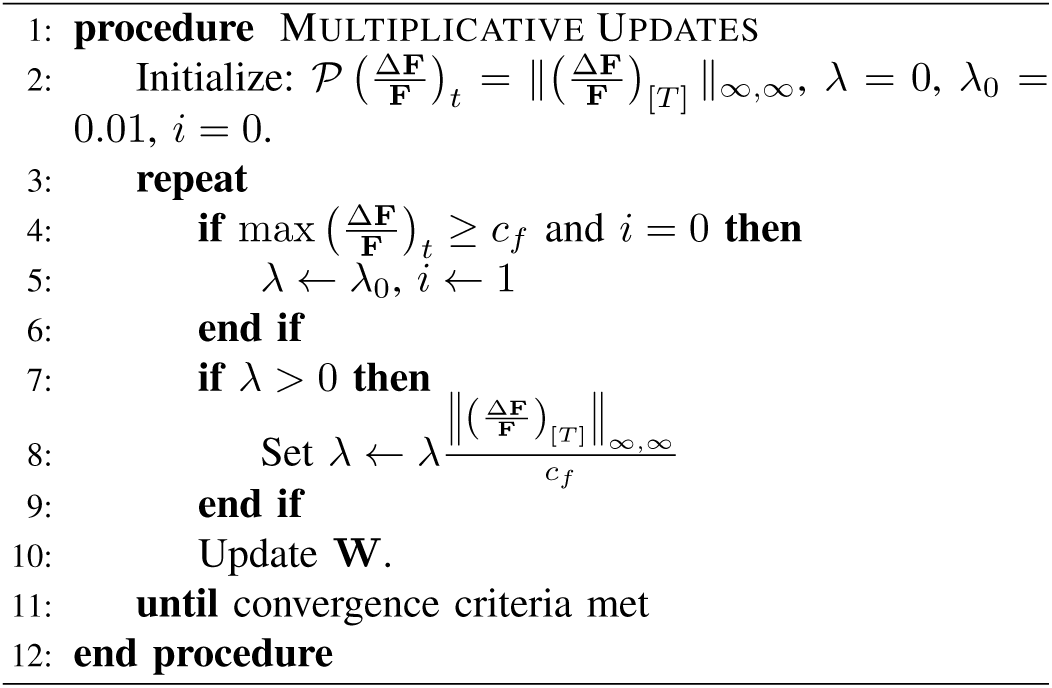

The main idea behind Algorithm 1 is that when the constraints are violated the complimentary slackness condition should be met for the optimal dual variable λ in Lagrangian form of the problem, meaning that the optimal solution should satisfy 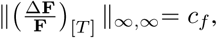 which is equivalent to finding a fixed point of updates for the dual (regularization) variable λ.

## V. Other Examples

### A. Dynamic Nonnegative Least Square (NLS)

The NLS problem can in general be formulated

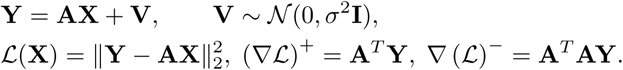

The most famous algorithm for solving the NLS problem is the active set method [4] which does not account for temporal dynamics in x_*t*_ or other forms of penalty. In these settings our update rules are very similar to the nonnegative deconvolution problem. A very useful example from the compressed sensing literature is the Multiple Measurement Vector (MMV) problem (without the positivity constraint) [27]. A commonly used penalty in this setup is the ║X║_2,1_ which enforces row sparsity.

### B. Dynamic Nonnegative Matrix Factorization (NMF)

The NMF problem is very similar to the NLS problem except that the matrix A is not known. In this case we can alternatively update our estimates of A and X [28].

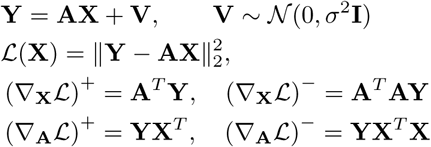

In the absence of penalization or dynamics we recover the multiplicative updates of [1]. Our update rules can also account for the dynamic case where

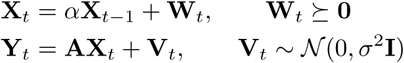

For example one can account for sparsely changing temporal factors by considering a Laplacian distribution on **W**_*t*_.

## VI. Extensions and Future Work

In this paper we considered convex optimization problems with nonnegativity constraints and provided unified multiplicative updates for them using the KKT conditions. These updates are easy to implement and parallelizable on a CPU. They do not require tuning of extra parameters such as the step size, exhibit fast convergence in practice and can account for temporal dynamics and smooth penalties without slowing down.

Although in the absence of convexity the KKT conditions no longer hold, we have empirically observed that our updates exhibit good performance when the problem has simple non-convexities. As an example one can model calcium saturation in the calcium deconvolution problem by adopting the calcium hill model given by 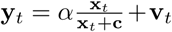 [29]. These observations suggest that suitable initializations result in convergence to a suitable local minimum. As another example one can combine the multiplicative updates with the IRLS algorithm [30] for 𝓁_*q*_, *q* < 1 minimization problems. The convergence of the IRLS algorithm was shown in the literature by showing an equivalence to a special case of the EM algorithm [31]. We applied this generalization to calcium imaging data using a nonconvex penalty.

Finally, the positivity constraint can easily be relaxed in the general form of the problems in two ways: First, any variable **X** can be decomposed into **X** = **X**^+^ – **X**^−^, where both **X**^+^ and **X**^−^ are positive. Second, generalized positivity and negativity could be defined with respect to the boundary of the convex set of feasible solutions, i.e. any point point inside/outside the feasibility set could be considered as positive/negative. Generalized positive and negative terms in the decompositions could be redefined similarly. Therefore by looking for a generalized positive fixed point of the gradient of the log-likelihood, the multiplicative updates can be generalized to a larger class of problems with not necessarily positivity constraints. We leave full details of these extensions and examples and their convergence properties to future work.

